# A human disease model of SARS-CoV-2-induced lung injury and immune responses with a microengineered organ chip

**DOI:** 10.1101/2020.07.20.211789

**Authors:** Min Zhang, Peng Wang, Ronghua Luo, Yaqing Wang, Zhongyu Li, Yaqiong Guo, Yulin Yao, Minghua Li, Tingting Tao, Wenwen Chen, Jianbao Han, Haitao Liu, Kangli Cui, Xu zhang, Yongtang Zheng, Jianhua Qin

## Abstract

Coronavirus disease 2019 (COVID-19) is a global pandemic caused by severe acute respiratory syndrome coronavirus 2 (SARS-CoV-2) that seriously endangers human health. There is an urgent need to build physiological relevant human models for deep understanding the complex organ-level disease processes and facilitating effective therapeutics for COVID-19. Here, we first report the use of microengineered alveolus chip to create a human disease model of lung injury and immune responses induced by native SARS-CoV-2 at organ-level. This biomimetic system is able to reconstitute the key features of human alveolar-capillary barrier by co-culture of alveolar epithelial and microvascular endothelial cells under microfluidic flow. The epithelial cells on chip showed higher susceptibility to SARS-CoV-2 infection than endothelial cells identified by viral spike protein expression. Transcriptional analysis showed distinct responses of two cell types to SARS-CoV-2 infection, including activated type I interferon (IFN-I) signaling pathway in epithelium and activated JAK-STAT signaling pathway in endothelium. Notably, in the presence of circulating immune cells, a series of alveolar pathological changes were observed, including the detachment of endothelial cells, recruitment of immune cells, and increased production of inflammatory cytokines (IL-6, IL-8, IL-1β and TNF-α). These new findings revealed a crucial role of immune cells in mediating lung injury and exacerbated inflammation. Treatment with antiviral compound remdesivir could suppress viral copy and alleviate the disruption of alveolar barrier integrity induced by viral infection. This bioengineered human organ chip system can closely mirror human-relevant lung pathogenesis and immune responses to SARS-CoV-2 infection, not possible by other *in vitro* models, which provides a promising and alternative platform for COVID-19 research and preclinical trials.

## Introduction

Coronavirus disease 2019 (COVID-19) pandemic broke out in late 2019 and quickly became a global epidemic [1–4]. Severe acute respiratory syndrome coronavirus 2 (SARS-CoV-2), the causative virus of COVID-19, has infected over ten million individuals up to July 2020 according to the report from World Health Organization (WHO), and the number of patients and deaths are increasing globally. COVID-19 patients exhibit a broad spectrum of disease progression and multiple clinical features including fever, dry cough, and ground-glass opacities [5, 6]. Human lung is the primary target for SARS-CoV-2 infection, which is characterized by the process ranging from mild syndrome to severe lung injury and multi-organ failure. Many severe cases of COVID-19 develop progressive respiratory failure, leading to death due to diffuse alveolar damage, inflammation and pneumonia [6–9]. Based on pathological features of COVID-19 by biopsy samples, inflammatory infiltration of mononuclear cells or lymphocytes could be observed in lung interstitial tissues or alveolar cavities [9, 10], Particularly, previous studies from severe patients have suggested that an excessive inflammatory cytokine storm induced by SARS-CoV-2 often result in aberrant immunopathology and lethal outcome. However, in-depth mechanism of pathogenesis of COVID-19 is still not clear.

Presently, SARS-CoV-2 infection is studied mostly relying on monolayer cultures of cell lines and primary tissue cells [11, 12], human organoids [13, 14], and animal models [15, 16]. However, all of these models have their limitations. For example, monolayer cell cultures are oversimplified and cannot exhibit the complex structure and functions of human organ-specific microenvironments *in vivo*. Human organoids (e.g., lung organoids) can provide multiple cell types and more complex tissue-specific functions, but they cannot model organ-level features of lung, such as tissue-tissue interfaces, blood flow, cross-talk between epithelium and endothelium, and immune cell-host responses, which are essential for the pathological progression of viral infection in pulmonary tissue *in vivo*, In addition, animal models of SARS-CoV-2 infection have been established for validating therapeutics of drugs or vaccines [15, 16]. However, given the species difference, their targeted organs or systemic responses to SARS-CoV-2 infection may be significant different from human individuals. Moreover, the expensive cost, time-consuming process and animal ethics should be seriously considered. As such, it is highly desirable to develop alternative preclinical models that can better reflect human-relevant organ pathophysiology and responses to accelerate SARS-CoV-2 research and candidate therapeutics for COVID-19.

Significant advances in bioengineered organs-on-chips technology have made it possible to reconstruct 3D human organotypic models *in vitro* by recapitulating the key functions of living organisms in microfluidic device, such as intestine, heart, liver, lung and kidney [17]. They are able to model human-relevant organ physiology, holding great potentials for disease studies and drug testing [18–21], Here, we first report the establishment of a microengineered human pulmonary model of SARS-CoV-2 infection on a chip that allows to recapitulate the human relevant lung pathophysiology and immune responses associated with COVID-19 *in vitro*. The biomimetic lung alveolar chip consisted of two channels (alveolar lumen and vascular channels) sandwiched with extracellular matrix (ECM)-coated porous membrane that enables to the co-culture of human alveolus epithelial cells, pulmonary microvascular endothelial cells, and immune cells under perfused media flow. This device can reproduce the critical features of human alveolar-capillary barrier by synergistically combining human epithelium-endothelium interactions, 3D ECM and mechanical fluid cues. Upon SARS-CoV-2 infection, we systematically analyzed the responses of distinct cell types to virus by immunostaining and RNA-sequencing (RNA-seq) analysis. With addition of circulating immune cells in the vascular channel, we examined the pathological changes of epithelium-endothelium interface and inflammatory responses at virus post-infection, as well as the potential anti-viral therapeutics. This human-organ-chip offers a robust microsystem to study human-relevant organ responses to infectious virus, which are difficult to reproduce in other *in vitro* models. It provides a new strategy for COVID-19 research and future preclinical trials.

## Results

### 1. Characterization of microengineered human alveolus chip

Lung is the primary target organ in the progress of COVID-19. Human alveoli contained alveolar-capillary barrier is an important component unit of lung. Alveolar-capillary barrier is composed of alveolar epithelial and vascular endothelial cells interacting across a 3D ECM, which plays vital roles in gas exchange and prevention of external hazardous substances invasion or virus infection (Fig. 1A). Clinical and histopathological evidences suggested alveolar function is seriously damaged by SARS-CoV-2. In order to create the human lung infection model by SARS-CoV-2 *in vitro*, we initially design and construct a biomimetic human alveolus chip in a multilayer microfluidic device under dynamic culture conditions. The microfluidic device consists of two channels separated by a thin and porous PDMS membrane coated with ECM (Fig. 1B). The PDMS membrane allows to form bio-interface and is beneficial to the interactions of upper and bottom cell layers. The culture chamber permits the perfusion of media flow, which can facilitate nutrients exchange and waste exclusion.

**Figure 1.**
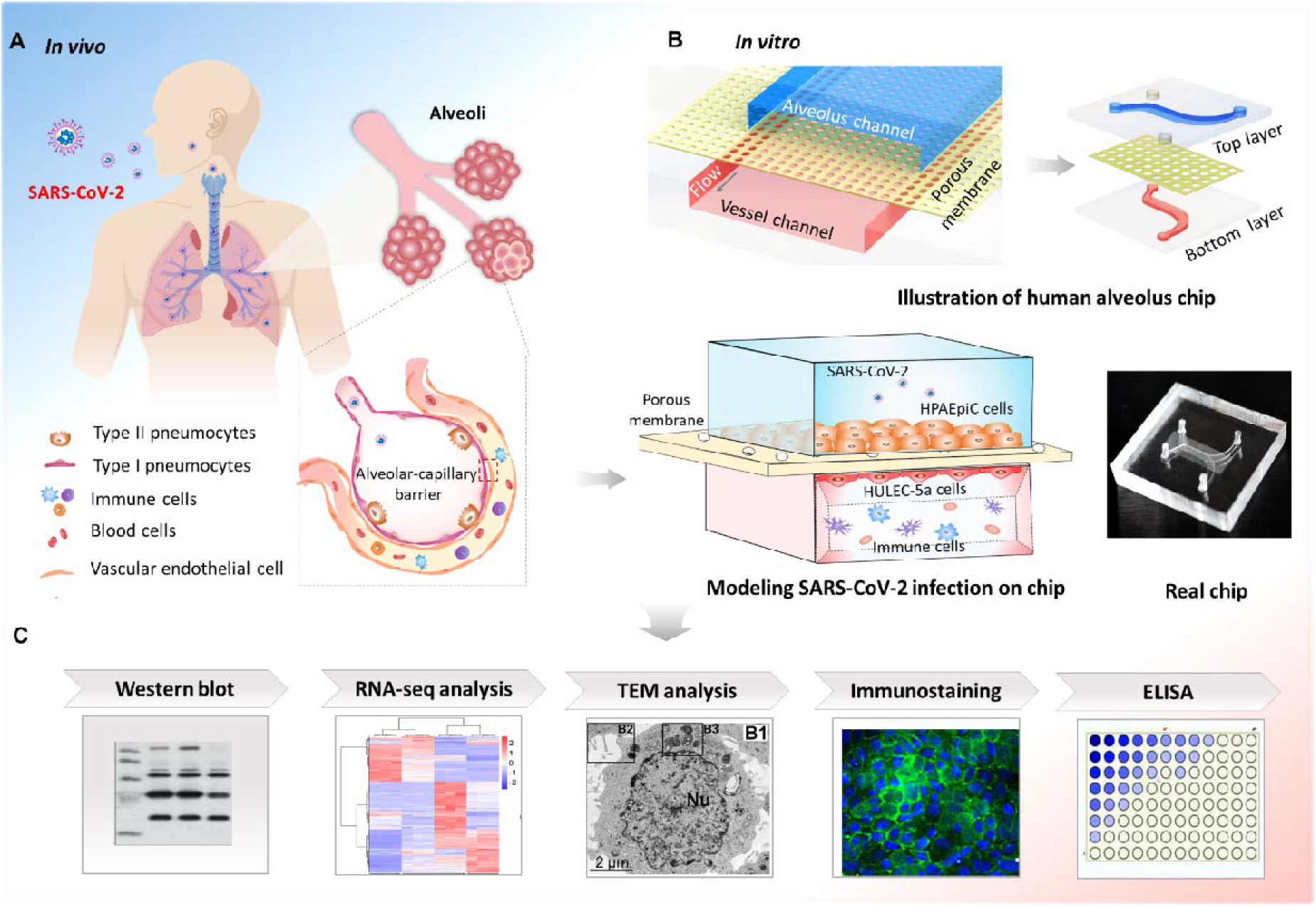
Schematic diagram of microengineered human alveolus chip infected by SARS-CoV-2. **(A)** Illustration of 3D human alveolar-capillary barrier *in vivo*. **(B)** The configuration of biomimetic human alveolus chip infected by SARS-CoV-2. The device consists of upper alveolar epithelial channel (blue) and lower pulmonary microvascular endothelial channel (red) separated by a porous PDMS membrane. The alveolar-capillary interface is formed by co-culture of alveolar epithelial cells (HPAEpiC) and pulmonary microvascular endothelial cells (HULEC-5a) on chip under fluid flow conditions. The established alveolus chip is exposed to SARS-CoV2 on the epithelial layer. Human immune cells are infused into the bottom vascular channel during the progression of virus infection. **(C)** The responses of human alveolus chip to SARS-CoV-2 are analyzed using different methods.

In this work, human alveolar epithelial type II cell (AT II) line (HPAEpiC) and lung microvasculature cell line (HULEC-5a) were seeded on the upper and lower side of porous membrane, separately. These two types of cells were cultured for 3 days until confluent into monolayers under continuous media flow (50 μl/h) in upper and bottom channels, thus forming an alveolus epithelium-endothelium tissue interface. The integrity of formed tissue barrier was assessed by the expression of adherent junction proteins in both human epithelium and endothelium. Immunostaining analysis showed that epithelial cells could form adherent junctions identified by E-cadherin, and endothelial cells formed conjunctions identified by VE-cadherin, respectively (supplementary Fig. S1). Furthermore, the integrity of barrier under different culture conditions was assessed by the diffusion rate of FITC-dextran between the two parallel channels (supplementary Fig. S2). The barrier permeability was lower under fluid flow than that in static cultures, indicating the important role of flow in maintaining the function and integrity of alveolar-capillary barrier.

### 2. SARS-CoV-2 infection in human alveolus chip

It has been reported that SARS-CoV-2 uses ACE2 as a host receptor for cellular entry, and transmembrane serine proteinase 2 (TMPRSS2) for Spike protein priming [12, 22–24]. Prior to create the SARS-CoV-2 infection model based on human alveolus chip, we sought to identify the susceptibility of alveolar epithelial cells to this virus. Initially, we examined the expression of ACE2 and TMPRSS2 proteins in HPAEpiC and HULEC-5a cells, respectively (Fig. 2A). The western blot data showed the positive expression of ACE2 and TMPRSS2 in both cell types [12, 25], and the higher ACE2 expression in HPAEpiC than HULEC-5a cells. HPAEpiC cells were then infected with SARS-CoV-2 at a MOI of 10 in monolayer cultures. At 3 days post-infection, more than 20 % Spike protein-positive cells were observed (Fig. 2B). To further examine the ultrastructure of SARS-CoV-2-infected cells, transmission electron microscope (TEM) analysis of mock- or SARS-CoV-2-infected HPAEpiC cells were performed (Fig. 2C, D). The TEM micrographs showed that mock cells exhibited primary AT II cell-like morphological characteristics, including a small cellular size with square or round shape (Fig. 2C1), microvilli on free surface (Fig. 2C2) and lamellar bodies within cell body (Fig. 2C3) [26, 27]. In infected cells, lots of viral particles were detected and distributed in clusters within cell bodies as shown in Fig. 2D, indicating the susceptibility of HPAEpiC cells to SARS-CoV-2 infection.

**Figure 2.**
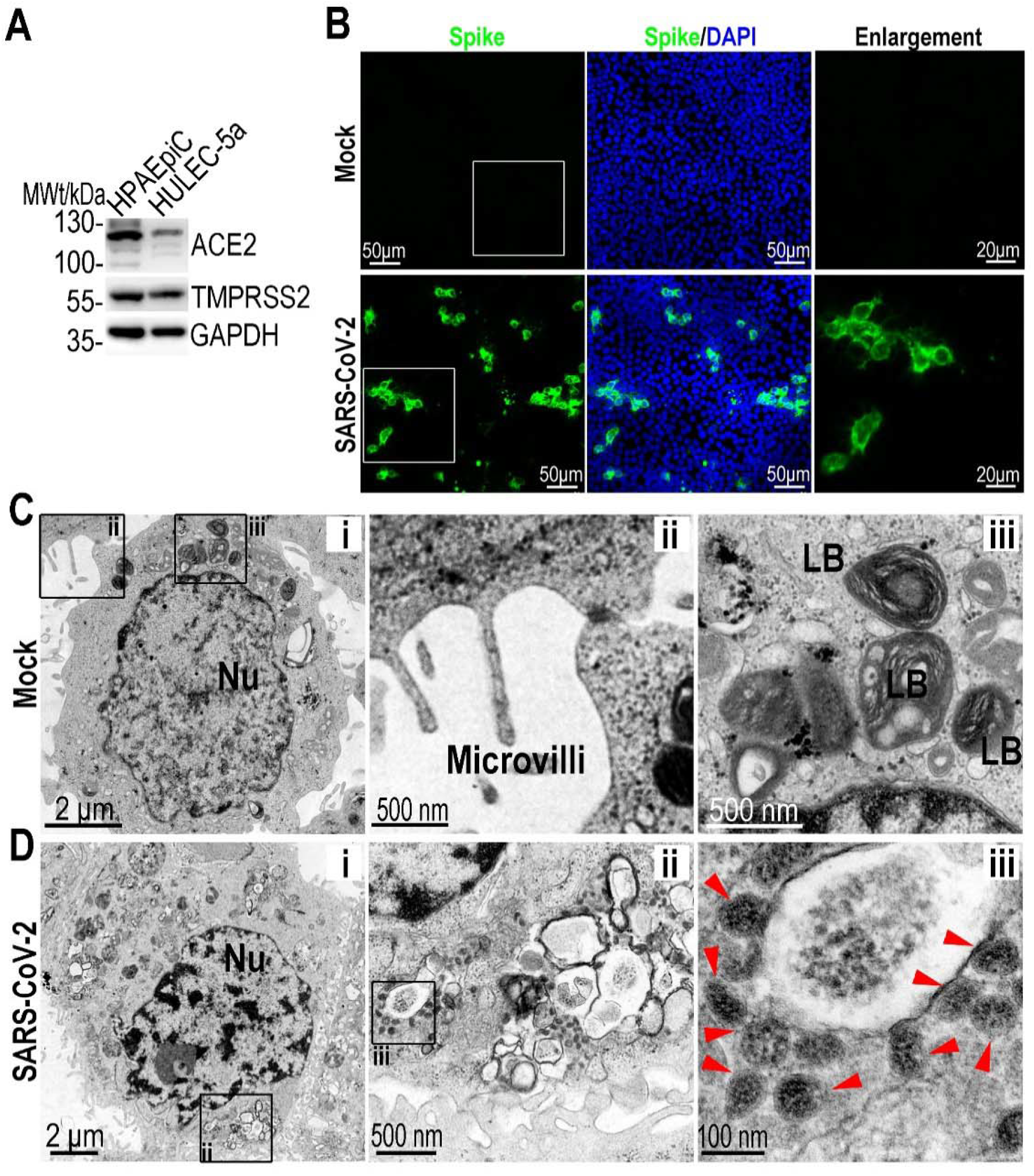
Examination of SARS-CoV-2 infection in HPAEpiC cells. **(A)** Analysis of ACE2 and TMPRSS2 proteins expression level in HPAEpiC and HULEC-5a cells by western blot. The results are representative blot from three experiments. GAPDH is served as a loading control. **(B)** Immunofluorescent images showed the viral Spike protein (Spike protein S1 subunit of SARS-CoV-2) staining in HPAEpiC cells at 72-hour post SARS-CoV-2 infection. **(C)** TEM micrographs of mock HPAEpiC cells without virus infection. (i) The overall image of HPAEpiC cell. (ii) The enlarged micrograph of microvilli on free surface of HPAEpiC cell. (iii) The enlarged micrograph of lamellar bodies (LB) within cell body. **(D)** TEM micrographs of SRAS-CoV-2-infected HPAEpiC cells. (i) The overall image of the infected cell. (ii) Clusters of viruses within cell body. (iii) Enlarged image of virus particles. Three independent experiments were performed (n=3).

To model alveolar infection by SARS-CoV-2, virus was inoculated into the upper alveolus channel of chip at a MOI of 10, and cell were cultured for 3 days. The predominated expression of spike protein was observed in epithelial cells, demonstrating the viral infection and massive replication in alveolar epithelium (Fig. 3A and B), but not in endothelial cells. In addition, there are not obvious changes in the organization of adherent junction proteins in HPAEpiC cells (E-cadherin) and HULEC-5a cells (VE-cadherin), as well as the confluent rate of epithelial cells and endothelial cells (Fig. 3C and D). These results suggested that SARS-CoV-2 can infect and replicate primarily in the alveolar epithelial cells, but not in endothelial cells.

**Figure 3.**
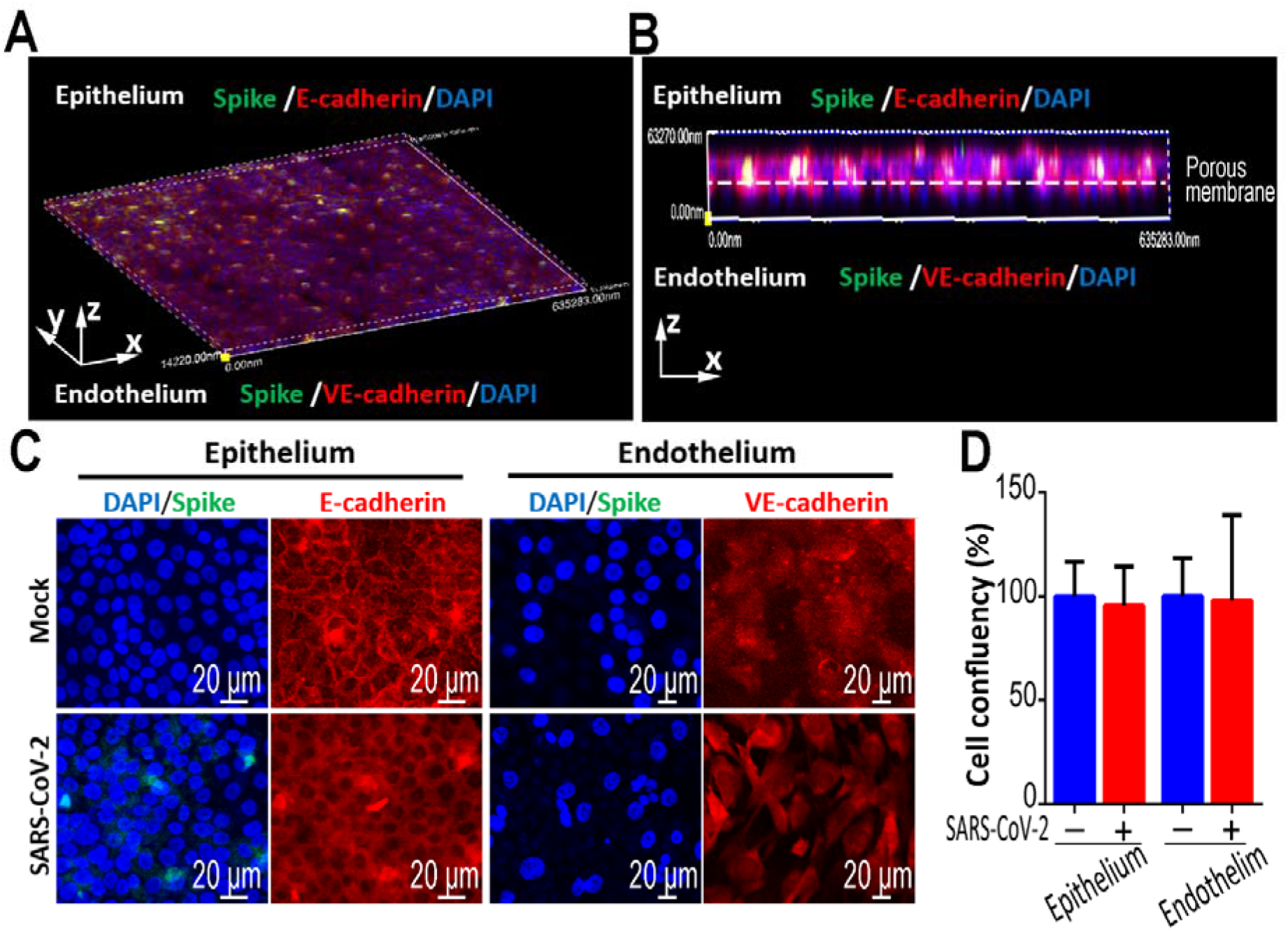
Characterization of infection and replication of SARS-CoV-2 in human alveolus chip. **(A)** 3D reconstructed confocal image of human alveolar-capillary-barrier at 3 days post-infection. **(B)** Side view of the formed alveolar barrier after viral infection. SARS-CoV-2 infection was identified in epithelium layer by viral Spike protein expression. **(C)** Confocal immunofluorescent micrographs showed the effects of SARS-CoV-2 infection (Spike protein) on the human epithelium (E-cadherin) and endothelium (VE-cadherin) of chip at 3 days post-infection. **(D)** Cell confluency of epithelium and endothelium on chip was examined with or without SARS-CoV-2 infection. Data were presented as mean ± SD. Three chips were quantified for each group.

### 3. Transcriptional analysis of host cells responses to SARS-CoV-2 infection

To gain a global understanding of transcriptional responses to SARS-CoV-2 infection, we performed RNA-seq analysis of HPAEpiC and HULEC-5a cells following virus infection in the alveolus chip. The ratio of virus-aligned reads over total reads in each sample was calculated to estimate the viral replication levels in these two cell types. The results showed that the ratio of viral reads was much higher in infected-HPAEpiC cells than HULEC-5a cells (Fig. 4A and B), which are consistent with the immunostaining analysis (Fig. 3B). It revealed that human alveolar epithelial cells were more permissive to SARS-CoV-2 infection than microvascular endothelial cells, similar to the histopathological findings from autopsy reports [28]. Hierarchical clustering analysis showed SARS-CoV-2 infection elicited broad transcriptional changes in both HPAEpiC and HULEC-5a cells (Fig. 4C). For identification of differentially expressed genes (DEGs), the cutoffs for the fold change and P value were set to 2.0 and 0.05, respectively. By combining the two data sets, we found the two cell types only shared 52 overlapping up-regulated DEGs (6.2% of total up-regulated DEGs) and only 43 overlapping down-regulated DEGs (5.5% of total down-regulated DEGs) (Fig. 4D). These results suggested that epithelial and endothelial cells displayed distinct transcriptome responses after SARS-CoV-2 infection.

**Figure 4.**
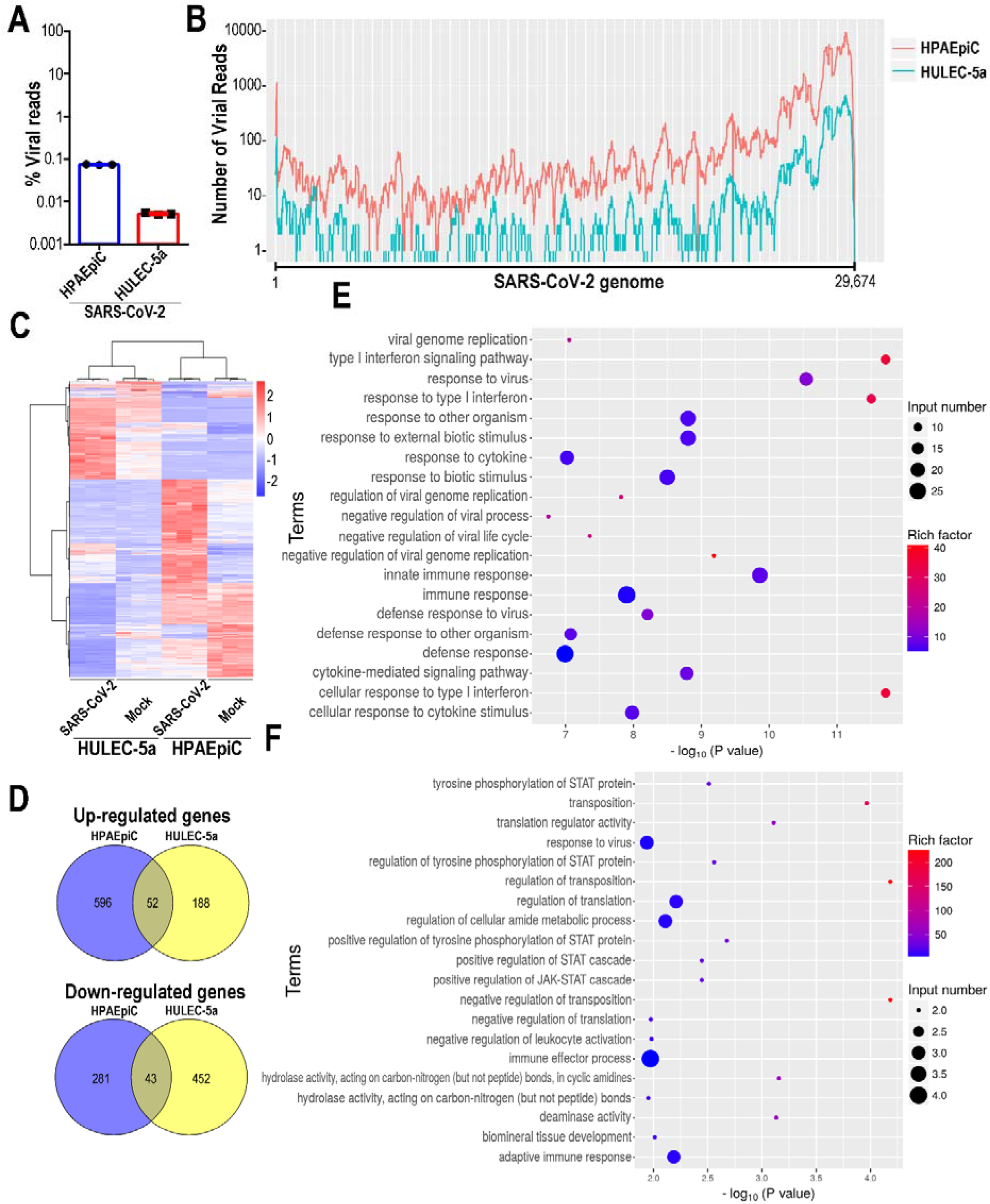
Transcriptional analysis of HPAEpiC cells and HULEC-5a cells to SARS-CoV-2 infection in human alveolus chip. **(A)** Viral replication levels in HPAEpiC and HULEC-5a cells. The ratio of virus-aligned reads over total reads is indicated for the viral replication level for each sample. Three independent experiments were performed (n=3). **(B)** Read coverage of viral reads along the SARS-CoV-2 genome for the infected-HPAEpiC cells or HULEC-5a cells. The graph indicated the viral reads number per position of the viral genome in the infected-HPAEpiC cells or HULEC-5a cells. This graph is representative of three independent experiments. **(C)** Heatmaps depicting the transcriptomic changes of HPAEpiC cells and HULEC-5a cells upon SARS-CoV-2 infection in alveolus chip. **(D)** Venn diagrams depicting the differentially expressed genes shared or unique between each comparison. For the identification of differentially expressed genes (DEG), the cutoffs for the fold change and P value were set to 1.5 and 0.05, respectively. **(E)** Dotplot showing the enriched GO terms in HPAEpiC cells following SARS-CoV-2 infection. The color of the dots represents the rich factor and the size represents the input number for each GO term. **(F)** Dotplot showing the enriched GO terms in HULEC-5a cells following SARS-CoV-2 infection. The color of the dots represents the rich factor and the size represents the input number for each GO term.

In general, viral infection can trigger antiviral or immune responses in host cells. We then performed gene ontology (GO) enrichment analysis of DEPs related with immune responses, and tried to identify which host defense responses can be induced after SARS-CoV-2 infection. The GO enrichment results showed, SARS-CoV-2 infection induced broader immune responses and antiviral responses in HPAEpiC cells, including defense response to virus, type I interferon (IFN-I) signaling pathway and cytokine-mediated signaling pathway (Fig. 4E). While, in HULEC-5a cells, terms related with positive regulation of JAK-STAT cascade and adaptive immune response were enriched (Fig. 4F). Moreover, among up-regulated genes related with immune responses in HULEC-5a cells, we identified some chemokines, including CCL15-CCL14, CCL15 and CCL23 (supplementary Fig. S3). Collectively, these findings revealed SARS-CoV-2 had distinct effects alveolar epithelial cells and microvascular endothelial cells, including the susceptibility to virus, immune responses and activated signaling pathway. This indicated the diverse roles of alveolar epithelial and endothelial cells involved in the pathogenesis of COVID-19.

### 4. Immune responses in human alveolus chip following SARS-CoV-2 infection

The accumulation and extensive infiltration of immune cells observed in the lungs may contribute significantly to the pathogenesis in patients infected with respiratory viruses [29]. We next explored the roles of human circulating immune cells in the alveolar pathological process after SARS-CoV-2 infection. In this study, PBMCs were isolated from healthy human blood and infused into the lower vascular channel of chip. SARS-CoV-2 was then inoculated into the upper channel at a MOI of 10 and incubated for 2 days. The infection of SARS-CoV-2 was identified in alveolar epithelial cells by Spike protein in the presence of PBMCs (Fig. 5A). Strikingly, in the presence of PBMCs, the distributions of both E-cadherin and VE-cadherin expressions in human epithelial and endothelial cells were intensely disrupted after virus infection (Fig. 5A). The obvious disruption of alveolar barrier function might be due to the involvement of PBMCs during the progress of viral infection. Furthermore, the cell confluency of human endothelium and epithelium after SARS-CoV-2 infection were examined on chip with the addition of PBMCs in vascular channel. Notably, the addition of PBMCs caused a significant decrease of endothelial cell confluency from 90.31±26.91% to 45.92± 8.52% (Fig.5B and C), however, no detectable changes of cell confluency were observed in epithelial cells. The data reflected the detachment and dysfunction of vascular endothelial cells in the presence of PBMCs in vascular channel. These results above revealed the crucial role of circulating immune cells in mediating the damage of alveolar barrier function after SARS-CoV-2 infection.

**Figure 5.**
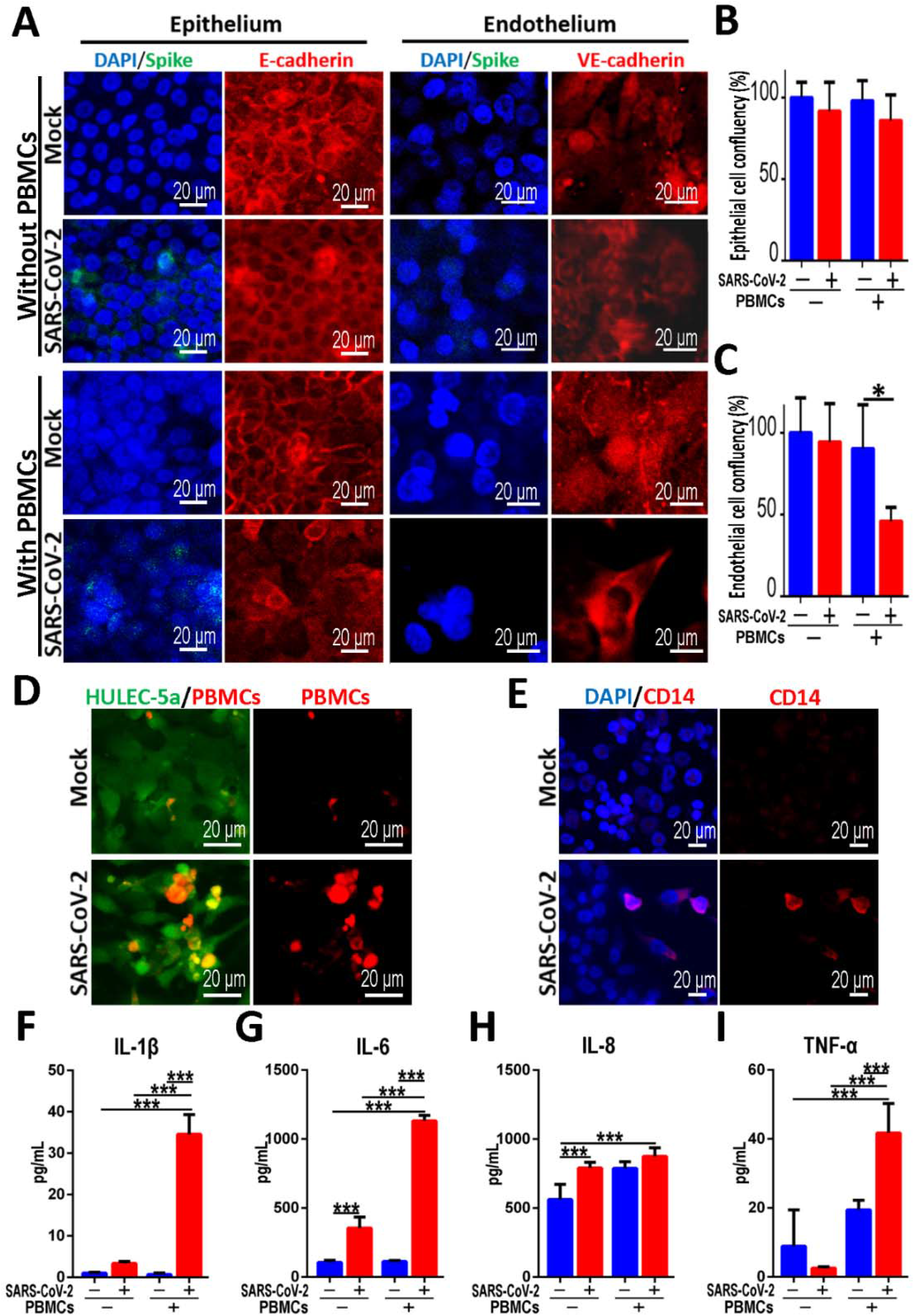
Distinctive responses of co-cultured epithelium and endothelium to SARS-CoV-2 infection in the presence or absence of circulating immune cells. **(A)** Confocal image showed the effects of SARS-CoV-2 infection (Spike) on the epithelium (E-cadherin) and endothelium (VE-cadherin) with or without immune cells (PBMCs) at 2 days post-infection. **(B-C)** Quantifications of cell confluency following SARS-CoV-2 infection in the presence or absence of circulating immune cells (PBMCs). Three chips were quantified for each group. Data were analyzed using a one-way ANOVA with Bonferroni post-test (*: p<0.05). **(D)** Confocal immunofluorescent micrographs showed the recruitment and adhesion of PBMCs (Red) on the surface of HULEC-5a layer (Green) after SARS-CoV-2 infection. **(E)** Confocal immunofluorescent micrographs showed the recruitment and adhesion of CD 14+ monocytes (Red) on the surface of HULEC-5a cell layer (Green) after SARS-CoV-2 infection. **(F-G)** Quantitative analysis of released inflammation cytokines IL-1β (F), IL-6 (B), IL-8 (C), and TNF-α (E) from the culture medium in vascular channel under different conditions. Data were presented as mean ± SD. Data were analyzed using a one-way ANOVA with Bonferroni post-test (***: p<0.001). Six chips were quantified for each group.

As above, transcriptional analysis had identified several up-regulated chemokines in endothelial cells after viral infection, which may be correlated with the recruitment of immune cells and inflammatory response. Subsequently, we examined the behaviors of circulating immunocytes in the vascular channel of chip after viral infection. Strikingly, the recruitment and adhesion of PBMCs, especially CD14^+^ monocytes were obviously observed on the surface of endothelium layer in the infected chip (Fig. 5D and E), which was similar to the inflammatory cell infiltration as observed clinically [9, 30].

To further explore the possible inflammatory responses induced by SARS-CoV-2 in this alveolus chip, we examined the production of several pro-inflammatory cytokines in culture media from the vascular channel. At 2 days post-infection, the levels of IL-6 and IL-8 were significantly increased in the absence of PBMCs (Fig. 5G and H). Moreover, all four cytokines (IL-1β, IL-6, IL-8 and TNF-α) were markedly increased following SARS-CoV-2 infection in the presence of PBMCs (Fig. 5F-I). Especially, viral infection caused a 10-fold increase in the level of IL-1β and IL-6 (Fig. 5F and G). We also examined the inflammatory cytokines secretion in culture media from the epithelial layer, which showed similar results to that from the vascular channel (supplementary Fig. S4). These results indicated that SARS-CoV-2 infection induced the recruitment of PBMCs, and aggravated inflammatory response, which are consistent with clinical findings. It appears this human organ chip could reflect human relevant pathological changes and host immune response to SARS-CoV-2, which are difficult to replicate in existing *in vitro* models.

### 5. Assessment of potential anti-viral therapeutics of remdesivir

To explore the potential therapeutics against SARS-CoV-2, we treated the virus infected human alveolus chip with remdesivir. Remdesivir is recognized as a promising antiviral compound against many RNA viruses (e.g., SARS, MERS-CoV), including SARS-CoV-2 [31–33]. Recent clinical trials showed remdesivir can shorten the disease course and reduce mortality of severe COVID-19 patients, and it has been approved by FDA in US [34]. In this study, an indicated dose of remdesivir (1μM) was added into the monolayer culture of HPAEpiC cells at 1h post-infection of SARS-CoV-2. After administration for 3 days, the culture supernatants were collected for virus titers determination by qRT-PCR. A marked decrease of virus titers was detected in the infected-HPAEpiC cells following remdesivir treatment (Fig. 6A). Furthermore, we tested the antiviral efficacy of remdesivir in the infected chip model with the addition of PBMCs in the vascular channel. As shown in Figure 6B and C, remdesivir treatment could restore the damage of epithelial layers and endothelial layer to some extent (Fig. 6B and C). These results indicated the potential role of remdesivir in suppressing SARS-CoV-2 replication and alleviating the virus-induced injury of alveolar-capillary barrier. It also suggested early administration of remdesivir might be helpful for disease control and alleviating lung injury for COVID-19 patients.

**Figure 6.**
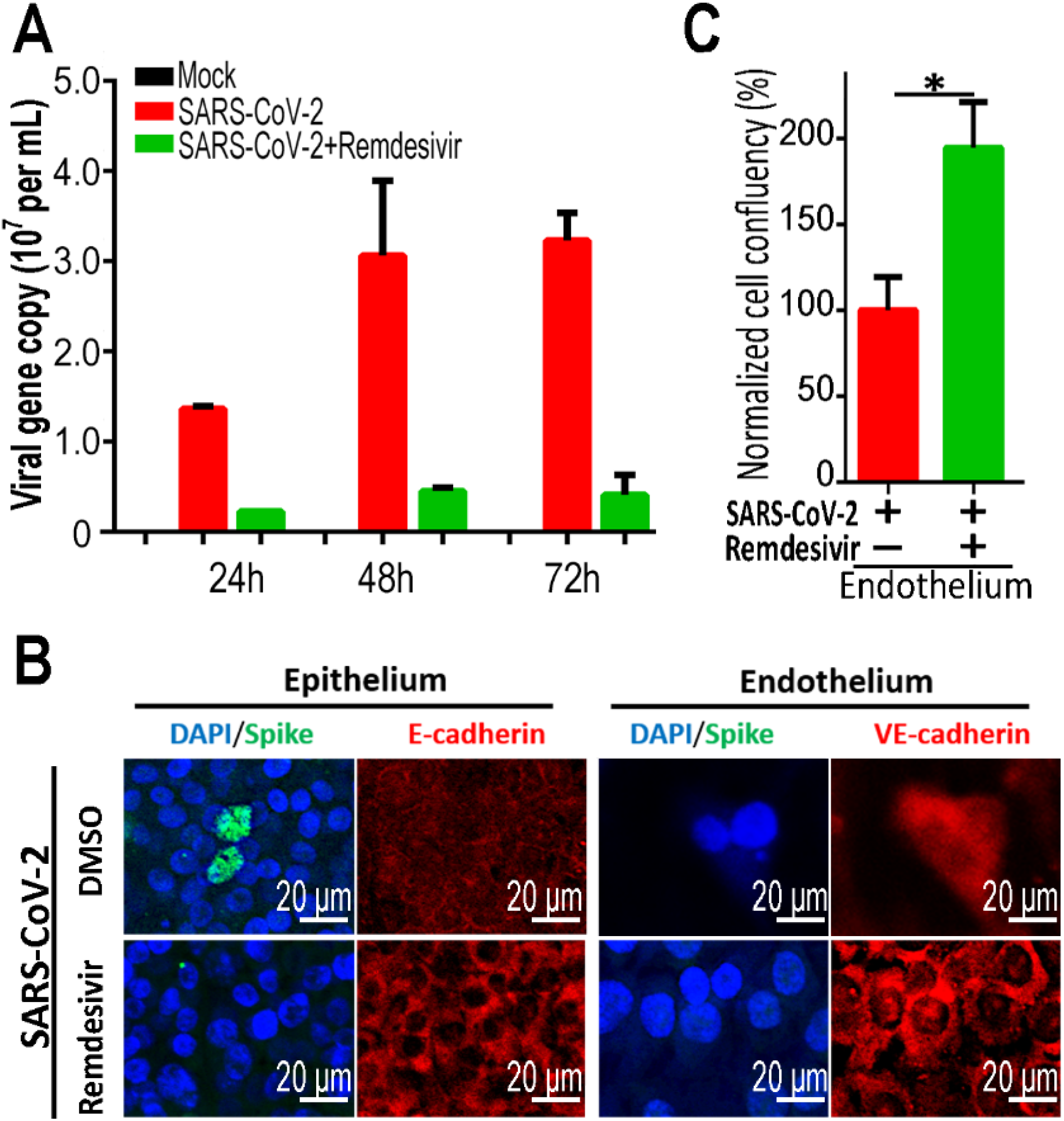
Evaluation of potential antiviral efficacy of remdesivir on the chip. **(A)** Culture supernatants were harvested at indicated time points following SARS-CoV-2 infection to examine the viral load using qRT-PCR for different groups. The average of two independent experiments is shown. Data were presented as mean ± SD. **(B)** Confocal immunofluorescent micrographs of epithelium (E-cadherin) and endothelium (VE-cadherin) of alveolus chip treated without or with 1 μM remdesivir 2 days post-infection. **(C)** Quantification for epithelial cell confluency of alveolus chip treated without or with 1 μM remdesivir at 2 days post-infection. Three chips were quantified for each group. Data were analyzed by Student’s t-test (*: p<0.05).

## Discussion

In this study, we created a microengineered human disease model of SARS-CoV-2 infection *in vitro* that allowed to recapitulate the pathophysiology and immune responses of alveoli at organ-level. The biomimetic human chip model closely recapitulated alveolar-capillary barrier injury and inflammatory responses following SARS-CoV-2 infection, such as viral replication in human alveolar epithelium, vascular dysfunction, recruitment of immune cells, and increased inflammatory cytokine release in a physiological-relevant manner (Fig. 7), which are not be readily modeled in existing *in vitro* model systems. Particularly, we found that human circulating immune cells play a key role in exacerbating inflammatory responses and the injury of alveolar-capillary-barrier induced by SARS-CoV-2. These findings provide new insight into the pathogenesis of SARS-CoV-2, in which virus-induced inflammatory responses and lung injury were possibly mediated by the complex and intrinsic cross-talk among epithelium-endothelium interface and immune cells.

**Figure 7.**
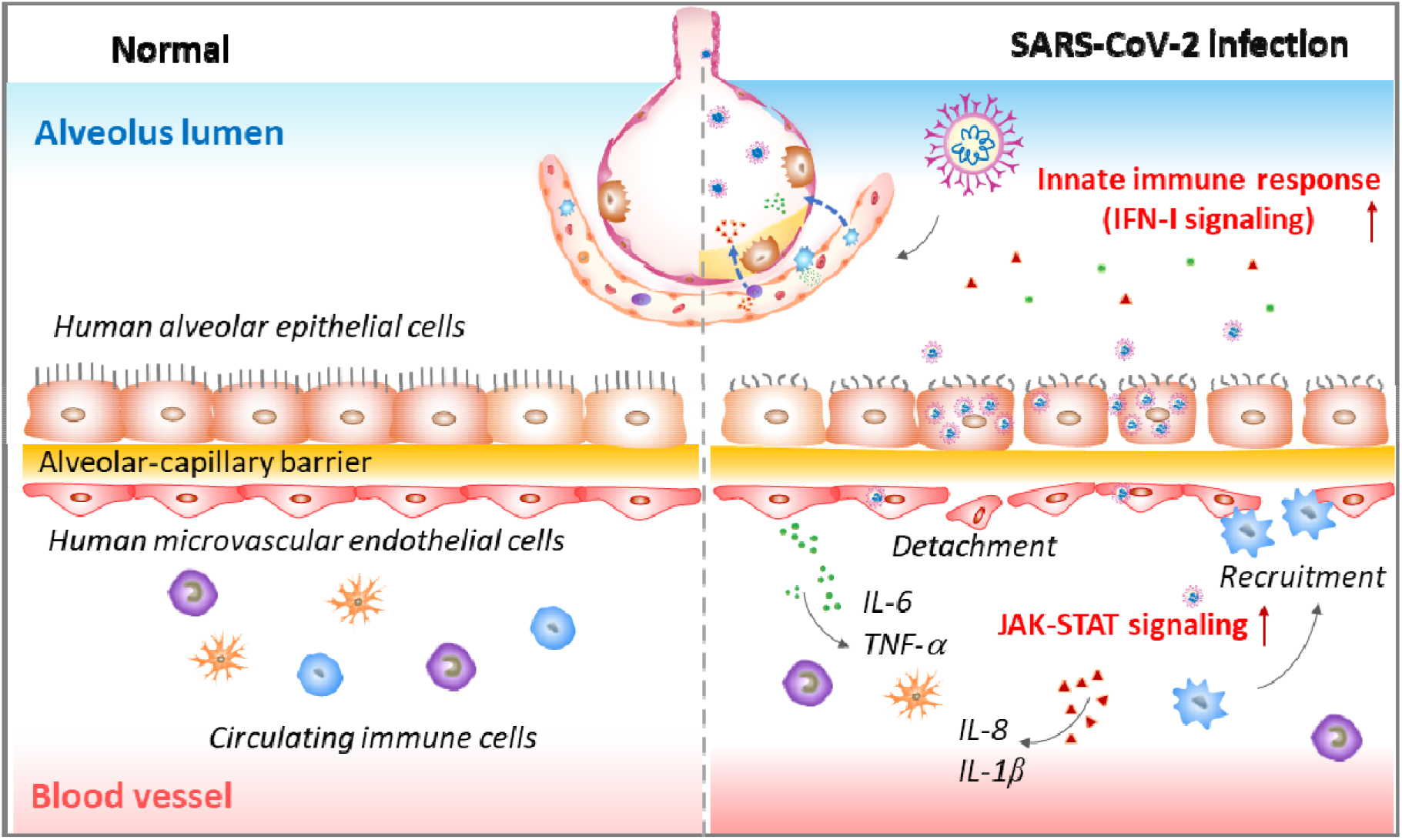
Proposed working model for SARS-CoV-2-induced lung injury and inflammatory responses based on human alveolus chip. Following SARS-CoV-2 exposure, virus particles invade the alveolar epithelium and endothelium, and replicate violently in human epithelial cells. The viral infection can activate the host antiviral defense or immune response, including the activation of innate immune responses (e.g. IFN-I signaling pathway) in epithelial cells and JAK-STAT signaling pathway in endothelial cells. The released cytokines or chemokines from infected cells can recruit circulating immune cells (such as CD14^+^ monocytes) to infected sites and initiate inflammatory responses. This process further exacerbates the disruption of alveolus-capillary barrier integrity, and leads to the lung injury.

As known, the AT II cells have been demonstrated as the primary infected target of SARS-CoV-2 by histopathological studies [35]. In this study, we found human alveolar epithelial cells were more susceptible to SARS-CoV-2 infection than endothelial cells as identified on the chip. Moreover, transcriptome analysis demonstrated a distinctive response of alveolar epithelial and endothelial cells to SARS-CoV-2 infection, in which epithelium exhibited much higher viral load than that of endothelium as observed on the co-cultured alveolus chip. This may partially explain why SARS-CoV-2 cannot be easily detected in the blood samples of COVID-19 patients [28]. In addition, compared with pulmonary microvascular endothelial cells, alveolar epithelial cells displayed broader and stronger innate immune responses and antiviral responses following viral infection, such as IFN-I signaling pathway. A recent study reported IFN-I responsive gene sets are up-regulated in lung tissues from severe COVID-19 deaths, which might be associated with exacerbated lung inflammation [36]. It appears our results are in agreement with the clinical results, indicating the feasibility of the lung infection model to reflect clinically relevant immune response in COVID-19. Meanwhile, the activation of JAK-STAT signaling pathway was observed in pulmonary microvascular endothelial cells after viral infection. It is well known that cytokines (e.g. IL-6) are enable to activate JAK-STAT pathway and further regulate different cellular and immune processes [37, 38]. Ruxolitinib, a JAK inhibitor could effectively relieve symptoms of patients with severe COVID-19 [39, 40]. Our findings suggested the potential therapeutic target for COVID-19 treatment by targeting JAK-STAT signaling pathway in microvascular endothelium.

Clinically, severe COVID-19 patients often display inflammatory cytokine storm that is associated with excessive immune responses, which may aggravate respiratory failure and cause multi-organ damage. Circulating cytokines, including IL-1β, IL-6, IL-8 and TNF-α are significantly elevated in patients with severe COVID-19 [41, 42]. Therefore, we detected these cytokines in the alveolus chip model following viral infection, and the results showed SARS-CoV-2 infection triggers secretions of these cytokines, and addition of circulating immune cells significantly further increases their level, accompany by recruitment of immune cells, and more severe damages of epithelium and endothelium. These data are relevant with pathological manifestations observed in severe patients with COVID-19 [43, 44]. Our findings revealed the key roles of immune cells in mediating alveolar injury, microvascular endothelial dysfunction and the excessive inflammatory response at organ-level.

One potential limitation of this work is the lack of human primary alveolar tissues that contains multicellular types of pneumocytes, such as alveolar epithelial type I and type II cells as existing *in vivo*. In addition, this model has yet to be tried for assessing more other drug candidates for COVID-19 therapeutics. Despite some limitations, the great value of the human alveolus chip is that it is capable to model human lung pathophysiology and study host-immune responses to respiratory viral infection at organ level. In combination with the existing cell-based models, this bioengineered lung infection model on chip may provide a complement to animal models for evaluating candidate drugs and repurposing approved drugs to face the crisis of SARS-CoV-2 epidemic.

Collectively, this work provides a proof-of-concept to establish a microengineered human disease model *in vitro* that enables to closely recapitulate lung pathophysiology and immune responses to native SARS-CoV-2 infection for the first time. Comprehensive analysis of this infection model revealed new insights into the pathogenesis associated with COVID-19, in which virus-induced inflammation is a major contributor lung injury with the involvement of circulating immune cells. This human organ chip system provided a synthetic strategy to rebuild human organs and analyze pathological responses at organ level by flexibly varying system parameters, which is opening up new avenues for COVID-19 research and drug development.

## Methods

### Device fabrication

The human alveolus chip device consisted of the upper and lower layers fabricated by casting Polydimethylsiloxane (PDMS) pre-polymer on molds prepared using conventional soft lithography procedures. 10:1 (wt/wt) PDMS base to curing agent (184 Silicone Elastomer, Dow Corning Corp) was polymerized to produce molded device with channels by thermal curing at 80 °C for 45 min. The top channel (1.5 mm wide × 0.25 mm high) and bottom channel (1.5 mm wide × 0.25 mm high) were used to form the alveolus lumen and the microvascular layer, respectively. The length of overlapping channels was 15 mm. The two channels were separated by a thin (~10 μm) through-hole PDMS membrane (5 μm pores) to construct tissue-tissue interfaces. The porous PDMS membranes were fabricated based on the glass templates and spin-coating method modified from the previous protocol. The membrane was sandwiched between the aligned upper and bottom channels of the device by oxygen plasma bonding for 30 s. Finally, the chip devices were sterilized in an autoclave and the porous member of the device was coated with rat tail collagen type I (200 μg/mL, Corning) on both sides at 37 °C for 48 h before cell seeding.

### Cell culture

African green monkey kidney epithelial Vero E6 cells (ATCC, no. 1586) were cultured in Dulbecco’s Modified Eagle’s Medium (DMEM, Gibco) supplemented with 10% Fetal Bovine Serum (FBS, Gibco). Immortalized human alveolar epithelial cells (HPAEpiC) were generated from Type II pneumocytes of human lung tissue (purchased form Sciencell Shanghai Corporation) and were maintained in RPMI 1640 medium (Gibco) supplemented with 10% FBS. Human lung microvasculature cell line HULEC-5a was purchased from Procell Corporation and were maintained in HULEC-5a growth medium (Procell, CM-0565). Human peripheral blood mononuclear cells were isolated from fresh human blood using Ficoll (Stem cell technologies) density centrifugation. Isolated PBMCs were resuspended in RPMI 1640 medium containing 10% FBS and 50 IU/mL IL-2 and used for adhesion assays on chip. All cells were cultured at 37 °C in a humidified atmosphere of 5% CO2.

To create the human alveolus chip, HULEC-5a cells (~ 1×10^5^ cells) were initially seeded on the bottom side of the collagen-coated porous PDMS membrane and allowed to attach on the membrane surface under static conditions for two hours. Subsequently, cells were washed with fresh medium to exclude/remove unattached ones. Then, HPAEpiC cells (~ 5×10^5^ cells) were seeded into the upper channel under static cultures. After cell attachment, constant media flows (50 μL/h) were applied in both the upper and bottom layers using peristaltic pump. The cells were grown to confluence for 3 days and the chips were maintained in an incubator with 5% CO2 at 37°C.

### Virus

A clinical isolate SARS-COV-2 strain 107 was obtained from Guangdong Provincial Center for Disease Control and Prevention, China, and propagated in Vero E6 cells. The virus titers were (infectious titers of virus) were determined by a TCID50 assay on Vero cells. All work involving live SARS-CoV-2 was performed in the Chinese Center for Disease Control and Prevention-approved BSL-3 laboratory of the Kunming Institute of Zoology, Chinese Academy of Science.

### SARS-CoV-2 Infections

HPAEpiC cells were seeded in 24-well plates (2×10^5^ cells per well) in RPMI 1640 medium containing 10% FBS. After seeding for 24h, cells were infected with SARS-CoV-2 at a MOI of 10. After one hour, cells were washed three times with PBS and kept in fresh medium for 3 days. At day 3 post-infection, cells were washed with PBS and then fixed with 4% paraformaldehyde (PFA) before analysis. The culture supernatant was collected for RNA extraction.

For SARS-CoV-2 infection in the human alveolus chip, the apical channel of chip device was infused with 30 μL of RPMI 1640 medium containing the indicated multiplicity of virus (MOI=10). After one hour of infection, cells were washed three times with PBS and kept in fresh medium. At day 3 post-infection, the apical and basal media were collected for analysis of released cytokines. The HPAEpiC and HULEC-5a cells cultured on chip were fixed for immunofluorescence analysis or lysed for RNA-seq data analysis, respectively.

### Permeability assay

The alveolar-capillary barrier permeability was assessed by detecting the FITC-dextran diffusion rate from the lower vessel layer to the upper alveolar channel. After a 3-day co-culture, the medium with FITC-dextran (40 kDa, 1 mg/mL) was then infused into the bottom channel of device. The media were collected from the upper channel at different time points (0 h, 1 h and 2 h) and the fluorescence intensity was measured using microplate system (ABI Vii 7).

### Western blot analysis

Protein samples were separated on 10% SDS-PAGE and then transferred onto 0.2 μm nitrocellulose membranes (GE Amersham). After being blocked with 5% BSA in TBST buffer containing 0.05% Tween-20, the membranes were probed with the anti-ACE2 antibody (1:1000 dilution), anti-TMPRSS2 antibody (1:1000 dilution) or anti-GAPDH antibody (1:2000 dilution) at 4ū overnight. The membranes were then probed with horseradish peroxidase (HRP)-conjugated secondary antibodies at room temperature for 1 hour at room temperature. Protein bands were detected by Prime Western Blotting Detection Reagent (GE life).

### Immunostaining

HPAEpiC cells cultured on well plate were washed with PBS and fixed with 4% PFA at 4□ overnight. Cells were then permeabilized with 0.2% Triton X-100 in PBS (PBST buffer) for 5 min and blocked with PBST buffer containing 5% normal goat serum for 30 minutes at room temperature. Antibodies were diluted with PBST buffer. Cells were stained with corresponding primary antibodies at 4°C overnight and with secondary antibodies (supplementary Table S1) at room temperature for 1 hour. After staining with secondary antibodies, cell nuclei were counterstained with DAPI.

For immunofluorescent imaging of alveolus chip, cells were washed with PBS through the upper and bottom channels and fixed with 4% PFA. The fixed tissues were subjected to immunofluorescence staining by the same procedure as described above. All images were acquired using a confocal fluorescent microscope system (Carl zeiss LSM880). Image processing was done using ImageJ (NIH).

### Analysis of inflammatory cytokines

To analyze the released cytokines, the media in the vessel channel were collected from each chip. The concentrations of IL-6, IL-8, IL-1β, and TNF-α were measured using the corresponding human ELISA kits (Biolegend, USA) according to the manufacturer’s instructions.

### Real-time quantitative PCR (qRT-PCR)

Virus titers were determined by viral RNA detection using qRT-PCR. The culture supernatant for each condition was harvested for RNA extraction using the HP Viral RNA Kit (Roche, Cat no. 11858882001) according to the manufacturer’s instructions. QRT-PCR was performed using One Step RT-PCR RNA direct real-time PCR master mix (TOYOBO, QRT-101A) in a PCR System (Applied Biosystem, ViiA™ 7). The primers of SARS-CoV-2 E gene were as follows: N-F: 5’-GGG GAA CTT CTC CTG CTA GAA T-3’; N-R: 5’-CAG ACA TTT TGC TCT CAA GCT G-3’; N-probe: 5’-TTG CTG CTG CTT GAC AGA TT-3’. PCR amplification was performed under the following conditions: 50□ for 10 min, and 95□ 1min, followed by 45 cycles consisting of 95 □ for 15 s, 60 □ for 45 s.

### Transmission electron microscopy

HPAEpiC cells were collected and fixed in 4% PFA (Electron Microscopy Sciences) and 2.5% glutaraldehyde (Electron Microscopy Sciences) at 4□ overnight. After washed with PBS for three times and fixed in 1% OsO4 buffer for 2 h, the samples were dehydrated with graded ethanol solutions, and embedded in Epon 812 resin (SPI). Ultrathin sections (70 nm) were stained with 2% uranyl acetate for 30 minutes and then lead citrate for 10 mins. Images were acquired with a JEM-1400 PLUS electron microscope.

### Remdesivir treatment

Stock solution of remdesivir (GS-5734) was prepared in DMSO. For HPAEpiC cell cultures on plate, after SARS-CoV-2 infection for 1 h, cells were treated with or without 1 μM remdesivir for 3 days and the supernatants were collected at distinct time points (24 h, 48 h and 72 h) for viral load analysis by qRT-PCR. For alveolus chip, cells were treated with or without 1 μM remdesivir for 2 days after SARS-CoV-2 infection in the presence of PBMCs. After 48 h, the cells were fixed for immunofluorescence analysis.

### RNA extraction, library preparation and sequencing

HPAEpiC cells and HULEC-5a cells were collected separately from the chips, and total RNAs were extracted from samples using TRIzol (Invitrogen) following the methods by Chomczynski et al. [45]. DNA digestion was carried out after RNA extraction by DNaseI. RNA quality was determined by examining A260/A280 with NanodropTM OneCspectrophotometer (Thermo Fisher Scientific Inc). RNA Integrity was confirmed by 1.5% agarose gel electrophoresis. Qualified RNAs were finally quantified by Qubit3.0 with QubitTM RNA Broad Range Assay kit (Life Technologies). 500 ng total RNAs were used for stranded RNA sequencing library preparation using KC-DigitalTM Stranded mRNA Library Prep Kit for Illumina^®^ (Catalog NO. DR08502, Wuhan SeqHealth Co., Ltd. China) following the manufacturer’s instruction. The kit eliminates duplication bias in PCR and sequencing steps, using unique molecular identifier (UMI) of 8 random bases to label the pre-amplified cDNA molecules. The library products corresponding to 200-500 bps were enriched, quantified and finally sequenced on Hiseq X 10 sequencer (Illumina).

### RNA-seq data analysis

Raw sequencing data was first filtered by Trimmomatic (version 0.36), low-quality reads were discarded and the reads contaminated with adaptor sequences were trimmed. Clean Reads were further treated with in-house scripts to eliminate duplication bias introduced in library preparation and sequencing. In brief, clean reads were first clustered according to the UMI sequences, in which reads with the same UMI sequence were grouped into the same cluster, resulting in 65,536 clusters. Reads in the same cluster were compared to each other by pairwise alignment, and then reads with sequence identity over 95% were extracted to a new sub-cluster. After all sub-clusters were generated, multiple sequence alignment was performed to get one consensus sequence for each sub-cluster. After these steps, any errors and biases introduced by PCR amplification or sequencing were eliminated.

The de-duplicated consensus sequences were used for standard RNA-seq analysis. They were mapped to the reference genome of *Homo sapiens* from Ensembl database (ftp://ftp.ensembl.org/pub/release-87/fasta/homo_sapiens/dna/) using STAR software (version 2.5.3a) with default parameters. Reads mapped to the exon regions of each gene were counted by featureCounts (Subread-1.5.1; Bioconductor) and then RPKMs were calculated. Genes differentially expressed between groups were identified using the edgeR package (version 3.12.1). A FDR corrected p-valueū cutoff of 0.05 and Fold-change cutoff of 2 were used to judge the statistical significance of gene expression differences. Gene ontology (GO) analysis and Kyoto encyclopedia of genes and genomes (KEGG) enrichment analysis for differentially expressed genes were both implemented by KOBAS software (version: 2.1.1) with a corrected P-value cutoff of 0.05 to judge statistically significant enrichment.

### Statistical analyses

Data were collected in Excel (Microsoft). Differences between two groups were analyzed using a Student’s t-test. Multiple group comparisons were performed using a one-way analysis of variance (ANOVA) followed by post-hoc tests. The bar graphs with error bars represent mean ± standard deviation (SD). Significance is indicated by asterisks: *, p < 0.05; **, p < 0.01; ***, p < 0.001.

## Data availability

All relevant data are available in the manuscript or supporting information.

## Acknowledgments

We thank Prof. Yonggang Yao (Kunming Institute of Zoology, CAS) for his strong support on this work. We thank Prof. Shiyong Liu (University of Science and Technology of China) for providing remdesivir compound. We thank Ms. Yingqi Guo (Kunming Institute of Zoology, CAS) for help with the preparation of samples for electron microscopy analysis.

## Supplementary figures

**Figure S1.**
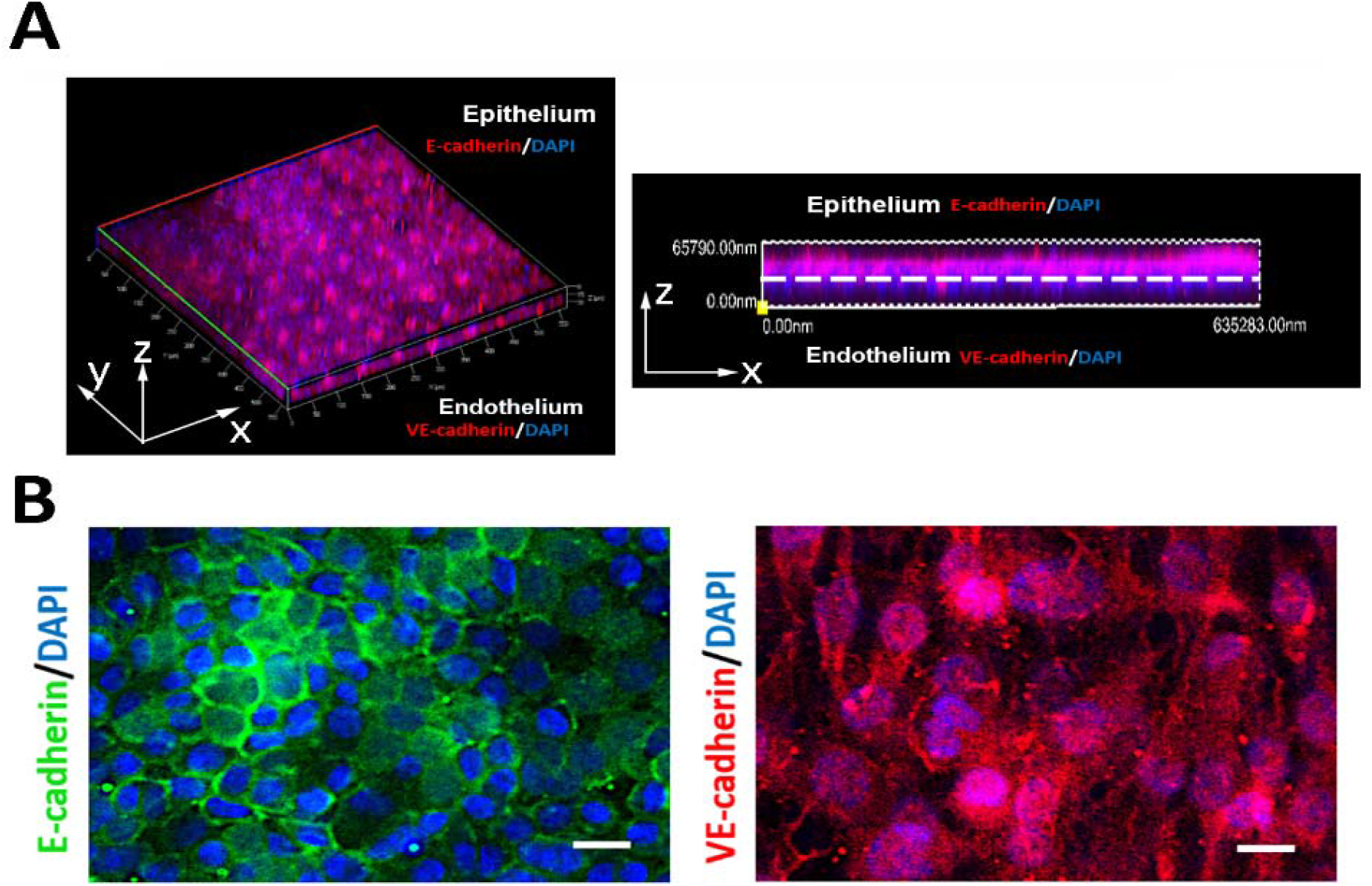
(A) 3D reconstructed confocal images of the formed human alveolar-capillary-barrier on chip after 3-day co-culture under fluid flow conditions. (B) The formation of adherent junctions in alveolar layer (E-cadherin) and microvascular endothelium (VE-cadherin) on alveolus chip. Scale bars: 20 μm.

**Figure S2.**
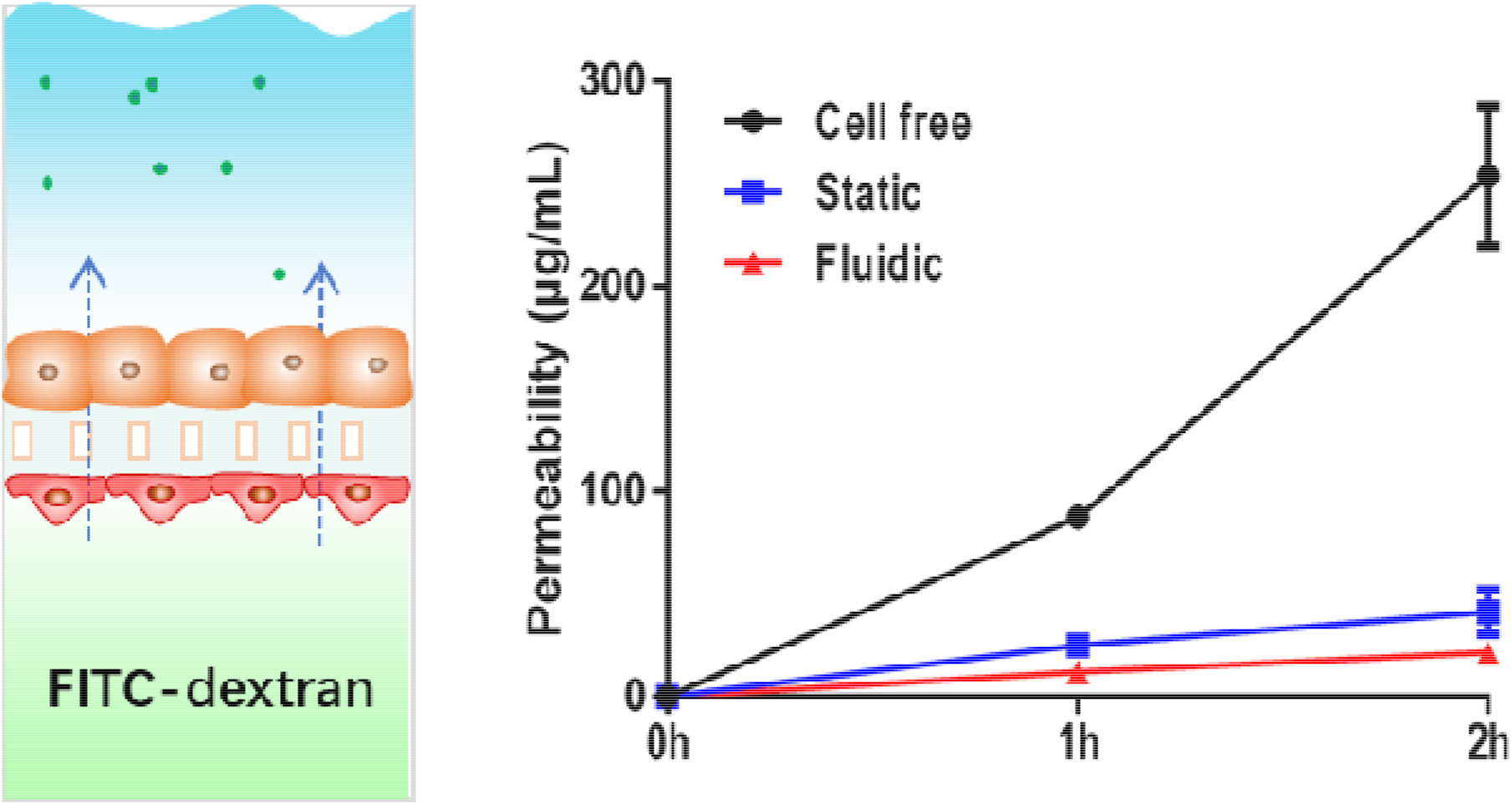
Permeability of the formed alveolar-capillary barrier within the alveolus chip under different culture conditions. After cell loading for 3 days, the medium with FITC-dextran (40 kDa, 1 mg/mL) was infused into the vascular channel of device. The media were collected from the alveolar channel at different time points (0, 1, 2 h) and the fluorescence intensity was measured to evaluate the permeability of the alveolar capillary barrier. Data were presented as mean ± SD. N=2.

**Figure S3.**
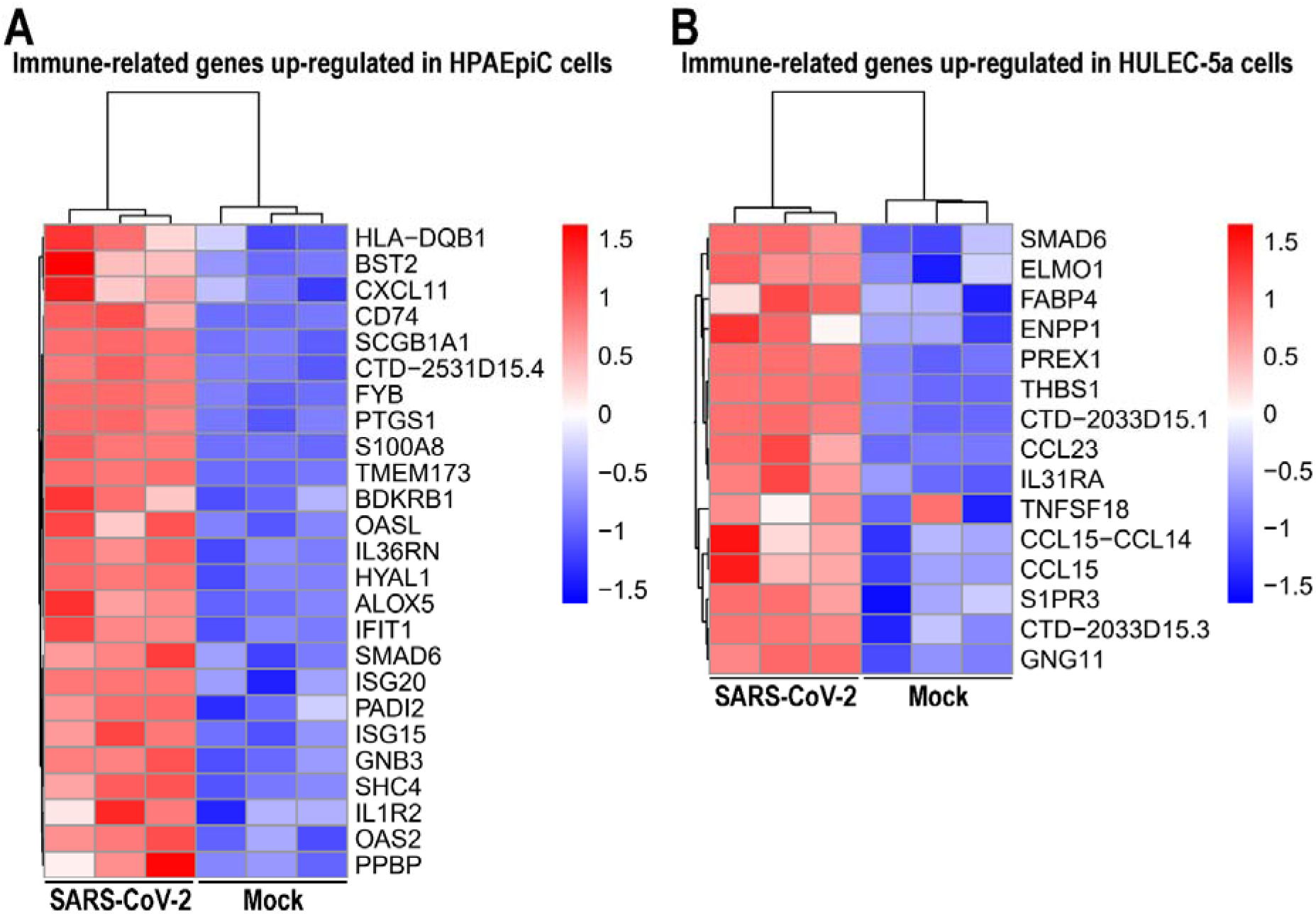
Heatmaps depicting the significantly upregulated genes related with immune responses upon SARS-CoV-2 infection in HPAEpiC (A) and HULEC-5a (B) cells, respectively. Colored bars represent Z-score of log2 transformed values.

**Figure S4.**
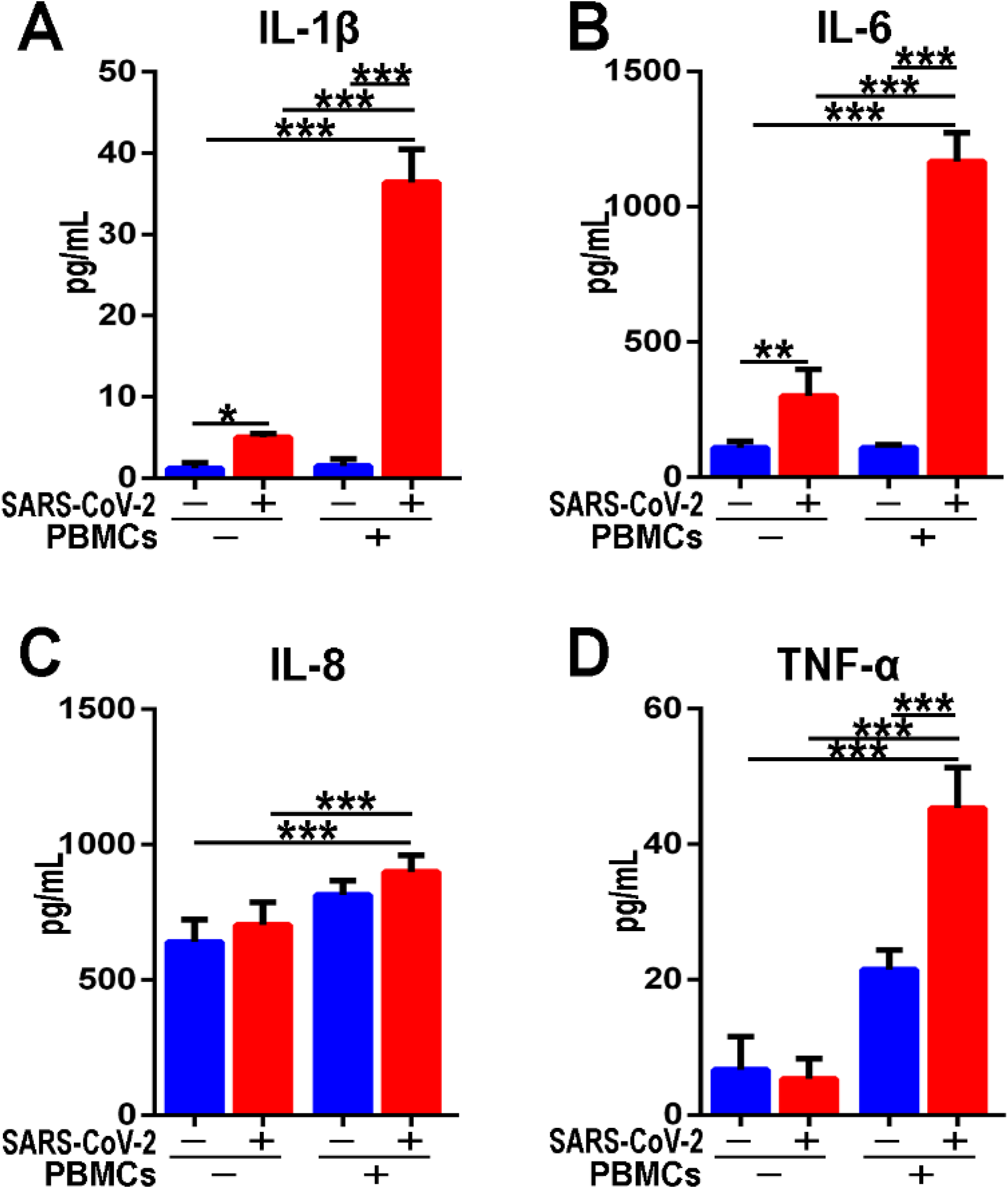
Examination of various cytokines production from epithelium effluence after SARS-CoV-2 infection in the presence or absence of PBMCs in the alveolus chip. Quantitative analysis of the secreted inflammatory cytokines (A) IL-1β, (B) IL-6, (C) IL-8, and (D) TNF-α in media from the alveolar layer. Data were presented as mean ± SD. Data were analyzed using a one-way ANOVA with Bonferroni post-test (***: p<0.001). Six chips were quantified for each group.

**Table S1.** List of antibodies used for western blot (WB) and immunofluorescence (IF).

| **Primary antibodies** | **Supplier** | **Cat #** | **Species** | **Purpose** | **Dilution** |
| --- | --- | --- | --- | --- | --- |
| ACE2 | Proteintech Group | 21115-1-AP | rabbit | WB | 1:1000 |
| TMPRSS2 | Proteintech Group | 14437-1-AP | rabbit | WB | 1:1000 |
| GAPDH | CWBIO | CW0100 | rabbit | WB | 1:5000 |
| E-caherin | Proteintech Group | 60335-1-Ig | mouse | IF | 1:200 |
| VE-cadherin | Proteintech Group | 66804-1-Ig | mouse | IF | 1:200 |
| CD14 | Abcam | Ab183322 | rabbit | IF | 1:200 |
| Spike | Sino Biological | 40150-R007 | rabbit | IF | 1:200 |

| **Secondary antibodies** | **Supplier** | **Cat #** | **Dilution** |
| --- | --- | --- | --- |
| Anti-Rabbit IgG(H+L), F(ab')<sub>2</sub> Fragment (Alexa Fluor® 488 Conjugate) | Cell Signaling | 4412 | 1:1000 |
| Anti-Rabbit IgG(H+L), F(ab')<sub>2</sub> Fragment (Alexa Fluor® 594 Conjugate) | Cell Signaling | 8889 | 1:1000 |
| Anti-mouse IgG (H+L), F(ab')<sub>2</sub> Fragment (Alexa Fluor® 594 Conjugate) | Cell Signaling | 8890 | 1:1000 |
| Anti-mouse IgG (H+L), F(ab')<sub>2</sub> Fragment (Alexa Fluor® 488 Conjugate) | Cell Signaling | 4408 | 1:1000 |

